# New genomic techniques, old divides: stakeholder attitudes towards new biotechnology regulation in the EU and UK

**DOI:** 10.1101/2023.06.04.543624

**Authors:** Jonathan Menary, Sebastian S. Fuller

**Affiliations:** Health Systems Collaborative, Centre for Tropical Medicine and Global Health, University of Oxford, Oxford, OX10SY, United Kingdom

## Abstract

The European Union and United Kingdom are in the process of establishing new regulation regarding the use of new genomic techniques in crop and animal breeding. As part of this process, consultations have been launched to understand the views of stakeholders and the wider public towards the use of new genomic techniques in plant and animal breeding. The responsible research and innovation framework emphasises the importance of dialogue between technology developers and stakeholders, including the public, but what are the opinions of stakeholders towards the regulation of NGTs in Europe and do they view these consultations as opportunities to engage with technology governance?

We conducted semi-structured interviews with experts from a range of agri-food stakeholder groups in the European Union and United Kingdom to understand current attitudes towards new biotechnology regulation, how they viewed the process of consultation in both places and what influence they felt they had in shaping regulations. We found that the discussion is similar in both EU and UK, with predictable and fixed opinions determined by attitudes towards the perceived risks associated with direct mutagenesis.

Both UK and EU consultations were considered to have the same weaknesses and stakeholders discussed a desire for more dialogic forms of engagement. We highlight several options for new forms of involvement in biotechnology regulation by exploring relevant responsible research and innovation literature.

## Introduction

The governance of new genomic techniques (NGTs) in plant breeding remains an issue of contention in the European Union and United Kingdom. In 2018, the European Court of Justice ruled that organisms obtained through directed mutagenesis would fall under the scope of existing legislation, namely the 2001/18/EC directive. In leaving the European Union in January 2020, the United Kingdom incorporated certain EU law into domestic law *–* including legislation covering the use of GM crops. In other parts of the world, crops developed through site-directed nuclease mutagenesis that do not include exogenous DNA (“SDN-1 and –2”) are not regulated as GMOs or exempted from existing regulations (van der Berg et al., 2021).

However, it has been suggested that the wording of the 2001 directive might allow for gene-edited crops to escape regulation in cases in which those crops could conceivably have been derived by conventional breeding (van der Meer et al., 2020). Due to this ambiguity and the wave of proposals to amend the directive following the Court’s ruling, the Council of the European Union asked the Commission to submit a report on NGTs, which was published in April 2021 (European Commission, 2021). Discussions have often centered on the status of point mutations and whether these should be excluded or exempt from existing regulation, given that these do not contain genetic admixture from non-sexually compatible species (Zimny & Eriksson, 2020). In the UK, ‘Brexit’ (the exit of the UK from the European Union) was cast by then prime minister Boris Johnson as an opportunity to change the law as it related to gene edited crops and animals (Stokstad, 2021). These initiatives come at time when agricultural and environmental policies are being reviewed in Europe through initiatives such as the *Farm to Fork Strategy* in the European Union and new, sustainable land management systems based on payment for public goods in the UK – biotechnology is noted as something that could play a role in meeting sustainability targets, but how newer techniques should be regulated is yet to be determined.

As part of the evidence-based policymaking strategy of the European Commission towards NGTs, a stakeholder consultation was launched in 2020 followed by a public consultation in 2022. The same is true in the UK, where a public consultation on gene-edited crops was launched by the Department for Environment, Food and Rural Affairs (Defra) in January 2021 before the results were published in September and the ‘Genetic Technology (Precision Breeding) Bill’ introduced in May 2022. The Bill is now law and requires developers to submit a ‘notification’ where new genomic techniques have been used to induce a mutation in a plant or animal, providing a route to market for these products that distinguishes them from first-generation GMOs (UK Parliament, 2023). (Due to the devolved nature of food and farming in the UK, the Bill only applies to England.) The consultations were presented as an opportunity for different views to be shared and have revealed ongoing divided opinion on more lenient regulation for gene-edit crops and animals (Poort et al., 2022). However, the extent to which these initiatives permitted viewpoints to be shared and feed into technology governance have been criticised; some organisations cast the language of the latest consultation as biased and chose not to participate (European Coordination Via Campesina, 2022); the UK public consultation was seen by some groups as ignoring the message provided by the exercise and criticised the weighting given to different groups’ opinions (A Bigger Conversation, 2021).

In recent years, there has been increased emphasis on providing the public with opportunities to engage and interact with innovations through collective decision making (Bruce & Bruce, 2019; see also Parker et al., 2014) and with biotechnology policy in particular (Bratlie et al., 2019; Kuiken et al., 2021; Stilgoe et al., 2013). Engagement has been promoted to improve acceptance of and trust in biotechnology and to build legitimacy in biotechnology policymaking (Gordon et al., 2021). This “deliberative turn” has its roots in conceptions of democratic forms of governance but can also function as a productive process that can, in itself, produce new knowledge for science and innovation (Macq et al., 2020). Questions remain as to who has the legitimacy to organise public discussion (Boëte, 2018) and how such discussions can influence the development of both the technology itself and the regulation governing its use. One framework for exploring these issues is responsible research and innovation (RRI).

### The PhotoBoost project and embedding RRI in crop improvement

We conducted this investigation as part of research for the PhotoBoost project. The Project is an international consortium funded by the European Commission Horizon 2020 programme to improve the photosynthetic performance of potatoes and rice, two of the world’s most important staple crops. There have been a number of recent advances in the field through optimising responses to light intensity in soybean (de Souza et al., 2022; Kromdijk et al., 2016). Heat and drought tolerance and water use efficiency are also benefits of improved photosynthetic performance (Baslam et al., 2020).

The quickest and most transformative options for improving photosynthetic performance rely on crop biotechnology (Long et al., 2006; Simkin et al., 2019). Due to a history of resistance to crops developed through the use of first-generation, transgenic techniques, there is a parallel need to integrate the upstream scientific understanding of photosynthesis with downstream (i.e. consumer and farmer) concerns and impact pathways (Kohli et al., 2020).

The RRI framework is a cornerstone of the Horizon 2020 funding programme. RRI emphasises the importance of strengthening public trust and understanding of science (Burget et al. 2017) and was developed in part as a response to the societal pushback, in Europe, to GM crops (Asveld et al., 2015). There is general agreement that responsible innovation should anticipate societal needs and be responsive to “changes in ethical, social and environmental impacts as a research programme develops” – this should include two-way consultation with the public and stakeholders (de Saille, 2015).

NGTs present an interesting challenge for the field, which is often associated with emergent technologies – such as synthetic biology or geoengineering (Stemerding et al., 2019; Stilgoe et al., 2013) – rather than technologies with a history of resistance like GMO crops (de Saille, 2015). Whether NGTs represent a break with older techniques (and constitute an emergent technology in their own right) is the subject of ongoing scientific, legal and social debate (see van der Meer et al., 2020; Zimny et al., 2019). However, Macnaghten (2016) argues that if RRI is to prove successful in aligning innovation with societal needs and values, it must be able to shape existing technological trajectories, such as GMOS, as well as those “in the making”. This can prove difficult given that entrenched power and influence, incumbent interests and technological “lock-in” serve to prevent societal shaping as technologies mature, even for new forms of biotechnology (Menary et al., 2020). PhotoBoost relies on newer (NGTs) and older (first-generation) biotechnology for crop improvement, so a dedicated work package was developed to explore stakeholder and public opinions towards biotechnology as it relates to improving photosynthesis.

Given recent interest in changing legislation related to the use of NGTs in the European Union and United Kingdom and the implementation of stakeholder engagement in the form of consultations, we sought to understand 1) the stance of different stakeholder groups towards the regulation of NGTs (and how these compare with first-generation biotechnology approaches), 2) how stakeholders viewed the consultatory process and 3) how stakeholders viewed their own and others’ influences on biotechnology policymaking. We also aimed to explore the extent to which responses differed between the EU and UK and how agri-food stakeholders viewed the aim of improving photosynthesis in the context of re-visiting biotechnology regulation. We contextualise our findings with reference to recent RRI and other relevant literatures.

### Methodology

We employed an applied qualitative approach that relied on semi-structured interviews with biotechnology and environmental policy experts, who were purposively sampled on the basis of their current professional position: 1) non-governmental and civil society organisation representatives, 2) industry and trade association representatives from the agri-food sphere and biotechnology research representatives. We sought a range of opinions from across the established ‘divide’ in positions on biotechnology regulation (from those in favour of stricter to more relaxed regulations) and sought a gender balance. We use the COREQ reporting criteria for qualitative research in presenting our findings (Tong et al., 2007). We gained ethical approval for the study from the *Social Sciences & Humanities Interdivisional Research Ethics Committee*, University of Oxford, reference R77406/RE001 in August 2021. The first author (JM) emailed a purposive sample of individuals and organisations who had responded to either the EU stakeholder consultation, or UK consultation (a list of respondents was published for each). Other participants who focus their research on biotechnology regulation in Europe were identified through literature searches. Participants who responded to an initial email invitation were provided with a participant information sheet (PIS), which outlines the aims and methods of the PhotoBoost project. If potential participants were satisfied with the information they received, a date for interview (via *Microsoft Teams*) was set and an online consent form link hosted at *JotForm*. Interviews were conducted by JM, who had no prior relationships with all but one interviewee. Participants were aware that the researcher was involved in PhotoBoost, which is funded by the European Union.

We also developed a ‘mapping exercise’ using online whiteboarding tool *Mural* for participants working in the European Union to visualise the agri-food policy-making environment and act as visual elicitation tool in order to prompt discussion around relative influence of different actors in that system (see Orr et al., 2020). Participants were shown the map and asked whether any notable system actors were absent – according to Glegg’s (2019) typology of visual research tools, this exercise helped us represent the data and enhance the quality and validity of our agri-food policy map.

We asked participants 1) how biotechnology should be regulated in the EU/UK, 2) what kind of information should inform that process and 3) if the consultation on new genomic techniques was the right approach. If participants were based in an EU country, they were shown the Mural conceptual map and asked 4) what organisations or entities are the most influential in determining agricultural biotechnology policy and 5) what pathways are used to influence policy. Participants in the UK were asked questions 1-3 but not consulted on the EU mapping exercise (i.e., questions 4 and 5).

Most interviews were audio and video recorded using Teams or audio recorded by *Dictaphone* when the use of Teams was not possible; one person was interviewed in person. All transcripts derived from recordings were pseudo-anonymised with identifying information redacted and original video/audio files stored on encrypted and password-protected *OneDrive for Business* drives. The transcripts were produced by a GDPR-compliant external company. Transcripts were checked and cleaned by JM prior to analysis.

The data were coded following the Framework approach by JM and SF (Ritchie et al., 2014). An initial coding structure was developed by each researcher independently from a sub-sample of three randomly-chosen transcripts and precise structure and themes were agreed upon. Subsequent indexing was carried out by JM using *NVivo 12*.

### Findings

We interviewed 15 people between September 2021 and November 2022 out of 52 (15/52, 29%) potential participants who were approached for interview: six of those contacted (6/52, 11%) felt that a colleague or acquaintance was more appropriate and passed on their details (all of whom agreed to interview); the remaining 31 (31/52, 60%) did not respond to emails. Saturation of themes occurred relatively quickly and was, as we discuss below, consistent across the EU and UK, which permitted limited the need for further sampling. Although there is no one-size-fits all approach to sampling (Fusch & Ness, 2015), Guest et al. (2006) find that saturation can occur at ∼12 interviews, with data variability dropping beyond that number.

The breakdown of participant’s roles is provided in Table 1 – we interviewed seven women and eight men (for reasons of confidentiality we do not provide specific job titles or list the organisations participants represent). Nine (9/15, 60%) belonged to organisations broadly in support of more lenient regulation of gene-edited crops, whilst five (5/15, 33%) wanted to maintain the current regulatory framework that sees gene-edited crops governed as genetically-modified organisms. One (1/15, 7%) stated neutrality on this issue.

**Table 1.**
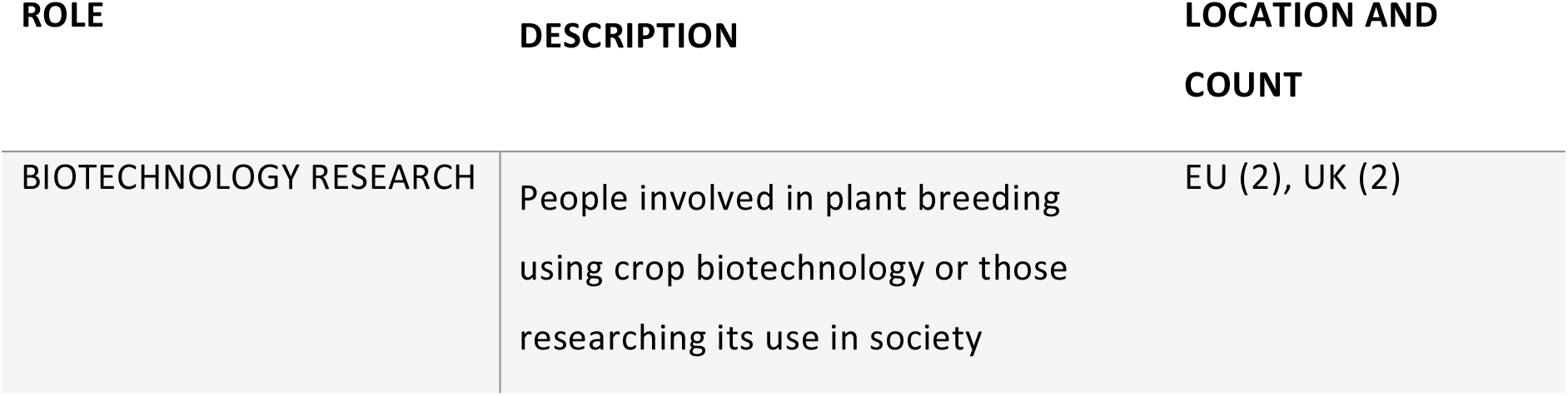
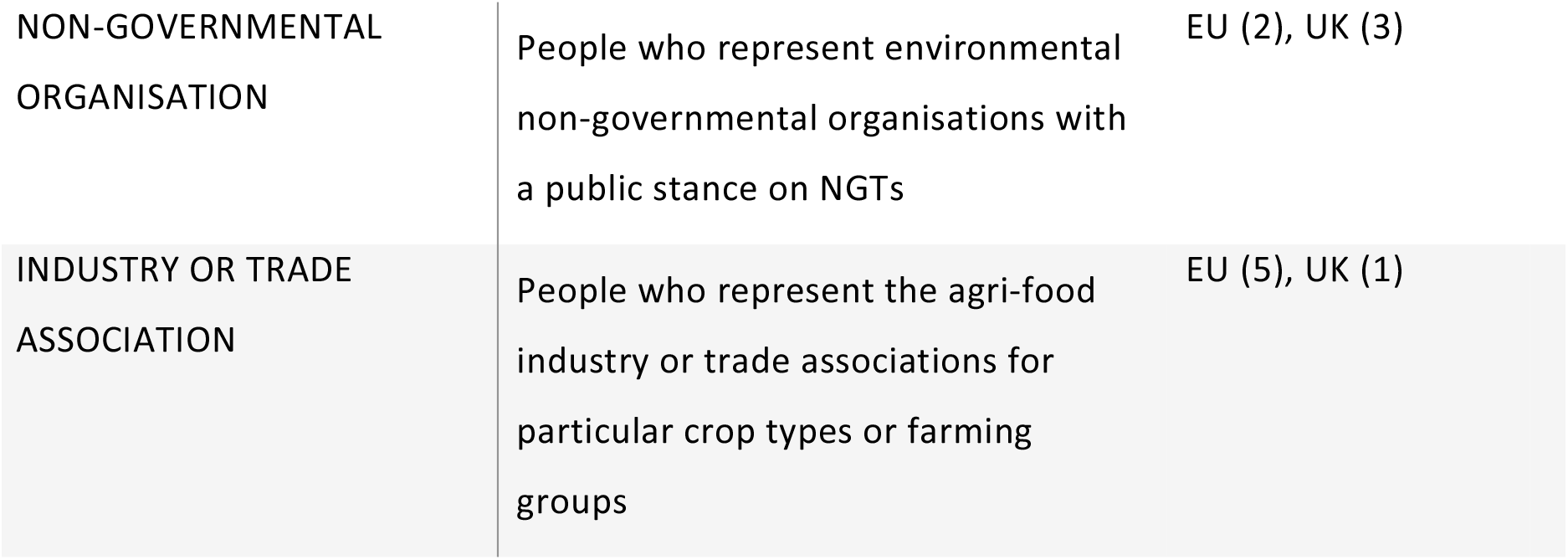
Participant role and professional location

| ROLE | DESCRIPTION | LOCATION AND COUNT |
| --- | --- | --- |
| BIOTECHNOLOGY RESEARCH | People involved in plant breeding using crop biotechnology or those researching its use in society | EU (2), UK (2) |
| NON-GOVERNMENTAL ORGANISATION | People who represent environmental non-governmental organisations with a public stance on NGTs | EU (2), UK (3) |
| INDUSTRY OR TRADE ASSOCIATION | People who represent the agri-food industry or trade associations for particular crop types or farming groups | EU (5), UK (1) |

A number of key preliminary themes were identified concerning the regulation of NGTs from our initial interviews.

#### 2.2.1 Biological lines in the sand

Participants’ views on how best to govern the use of NGTs were commonly rooted in biological risk. For those against softening regulatory processes for gene-edited crops, direct modification of the plant genome was paramount:

> *It’s not about whether it contains foreign genes or not, it’s this direct modification of the genome, which, frankly, is imperfect. It’s an imperfect science. People don’t really understand all the goings on in the genomes so how can you manipulate it directly and expect a sort of predictable outcome. – Non-governmental organisation (NGO) representative 1*

In particular, the unknown impacts of off-target genetic effects were invoked to cast doubt on claims about the specificity and relatively benign nature of single point mutations. These individuals called for *process-based* assessment of crops derived from NGTs. Conversely, participants calling for product-based regulation of such crops leaned on the fact that NGTs lead to changes indistinguishable from those derived from conventional breeding:

> *… introducing a gene that is crossable at the direct place where there’s inter-crossable species. And, [once] there’s no CRISPR or any other genome-editing tool left in the plant, I think it should be considered as natural and not as a GMO. And this is where… the line should be drawn. So, cisgenesis should not be regulated as a GMO. – Biotechnology research representative 2*

Other considerations – such as attitudes towards the risks and benefits of NGTs, detection and regulation – followed from participants’ position towards the biological basis of NGTs. Those who perceived risk in any direct modification of plant genomes, (e.g., NGO representatives), saw inherent risk in the use of crop biotechnology – those who viewed organisms modified by NGTs as no different from ‘natural’ equivalents saw no more risk than that associated with conventional breeding and tended to focus on the benefits of NGTs rather than risks. Few differences were attributable to the EU-UK divide in our sample but tracked rather more closely with participants’ organisations and occupations.

#### 2.2.2 Risks and benefits of biotechnology

The claim that crop biotechnology had in general not lived up to the promises made by its proponents was a common theme amongst those working for NGOs in both the EU and UK. The non-delivery on these promises was seen as part of the reason GM crops have been difficult to commercialise in Europe:

> *I think if the big promises would be fulfilled, then it would be very easy to go to any regulation process to any authorisation. Everybody would dare to use these plants. So, let’s go for it. Let’s just prove me wrong. Please, prove me wrong. – NGO representative 3*

And, because little distinction was drawn between NGTs and first-generation techniques by participants opposed to softening regulation, NGTs have inherited the legacy of those past failures, which they argued should be reflected in current policy:

> *Well GM was supposed to do all those things and it didn’t, so I am quite cynical about what [gene editing] actually will do. But the use of hypothetical potential as a reason why we should allow free rein is just extraordinary. – NGO representative 2*

The most cited risk of NGTs was unknown or unintended consequences of directed mutagenesis, both to the environment through genetic contamination and to human health through consumption of modified crops. Significant emphasis was placed on the current understanding of plant genetics as being insufficient to adequately appraise these unintended consequences.

Improved disease resistance, faster breeding times and heat stress and drought tolerance were cited as key benefits of NGTs by biotechnology researchers and industry representatives, as was reducing pesticide and fertiliser use. Improving the nutritional profile of crops and reducing levels of toxigenic compounds found in certain foods were also seen as key benefits. Some biotechnology researchers emphasised the risk of *not* using crop biotechnology: European competitiveness in plant breeding was invoked as in danger of falling behind other parts of the world, as was the ability to provide European farmers and consumers with improved varieties and food. A juxtaposition was also made by industry representatives between the success of COVID-19 vaccines and the perceived reluctance of the European community to trust in a similar branch of science. This group also saw the lack of acute food insecurity in Europe as contributing to a *laissez faire* attitude towards the need for new technology in agriculture.

The debate was characterised by NGO representatives as being about narrow “technological fixes” versus wider systemic change in the agri-food sector – change, which, in their view, meant much more systemic change in the food system (and society) rather than tweaking the existing model of agricultural production, which is how the goal of improving photosynthesis was cast:

> *“If it’s world hunger, people are not starving because photosynthesis is a bit rubbish. People are starving because they are poor and they’re not going to stop being poor because you’ve slightly improved the photosynthesis in their rice*… *it’s not as if we haven’t been having the debate for a long time that the kind of productivist regime, more and more calories*… *the issue isn’t production it’s actually access and nutrition*.*” – NGO Representative 2*

#### 2.2.3 Considerations for regulation and risk assessment

Numerous factors were considered by participants to be relevant to crop biotechnology regulation. The relevance of the 2001/18/EC directive, which provided the first definition for GMOs, was frequently discussed, particularly by those participants hoping to see more lenient regulation of SDN-1 techniques. Some suggested that the regulatory architecture of the European Union was sufficient without 2001/18/EC to deal with issues around novel food. According to this view, a product-rather than process-based form of regulation in which any mutagenic events are assessed on a case-by-case basis would make process-based legislation redundant. However, the most common view amongst industry associations we interviewed was to push for a technical amendment to the 2001/18/EC annexes, which includes exemptions to the directive. Conversely, NGO representatives we interviewed were either content with the regulation as it stood – providing gene-edited crops remained covered by the Directive – or could even be improved:

> *… we are pretty fine with the regulation like it is. We see gaps in there or still and maybe, especially when it comes to the authorisation process and the whole risk assessment. But, overall, we are fine with that and we think that genome editing should be regulated like the transgenic and cisgenesis tools. – NGO representative 3*

This included participants from UK-based NGOs, who saw the Government’s recent announcement regarding gene editing as a departure from the UK’s ‘inherited’ strict EU regulation on GMOs. When asked whether the wider agricultural industry sees a difference between newer and older biotechnology techniques, one participant noted that:

> *It has the same issues, the same debates, the same groups in favour and against, nothing has changed in that regard. – Biotechnology research representative 3*

Patenting and intellectual property (IP) were key concerns for NGO representatives, who saw the potential for easier patenting of gene-edited crops as furthering corporate control of food systems. The issues of traceability and product labelling were also important for many participants, although views on these issues, in contrast to discussions around product-versus process-based regulations, were less consistent amongst proponents of NGTs. The integrity of the Community Plant Variety Office (CPVO) rights system was considered critical for proponents of NGTs in the EU because breeders would need to know the provenance of certain varieties – or, more specifically, traits – for their own breeding programmes. There was concern from both industry and NGO representatives, for example, that non-labelled, gene-edited varieties might somehow threaten organic certification. One participant involved in plant breeding saw advantages in labelling food containing genetically-modified crops where the benefits of those modifications could also be promoted, whereas participants against relaxing regulation on gene-edited crops were more likely to view labelling as essential for consumer choice.

In the UK, NGO representatives feared that the permissive use of gene editing would “pull GM [crops] into that space”; a plant scientist worried that the promotion of gene editing as fundamentally different from first-generation transgenics – and therefore superior – had the effect of tarnishing what remained in their opinion a safe technology. Questions were raised over whether the UK would be able to export certain foodstuffs to the EU if the Precision Breeding Bill became law.

Whilst one of the often-promoted benefits of gene editing is single point mutation, NGO representatives were keen to point out that plant breeding programmes are rarely limited to one goal, with other changes introduced during the breeding process:

> *… they* [plant scientists] *do a lot of other things along the way. So, you know, it’s all very well saying ‘oh, we can just check for the thing we’ve added’, but you haven’t just added the thing that you’ve added. You’ve added a load of code either side of it, you’ve added a load of reagents to cause it to be entered into the cell. – NGO representative 2*

The number of discrete modifications made in the breeding process has a bearing on risk assessment, another issue that followed the division between the sampled groups. For most biotechnology research and industry representatives, a science-based risk assessment was not seen as controversial. A biotechnology research representative pointed to the inherently political nature of risk assessment and questioned “who” decides what factors are considered:

> *… these are all people who have a specific world view around the risk, they have a certain view of risk and of the value of biotech and science… so recognising that regulations and risk assessment are political spaces is the first important thing. – Biotechnology research representative 4*

Participants representing NGOs also questioned whether risk assessment adequately captured the “balance of interests” in European agriculture and promoted a wider form of assessment that included social and economic considerations.

#### 2.2.4 Views on consultations

When it came to the consultation processes employed the Commission and by the UK Government, our participants considered that the intent of a consultation process was correct and necessary. However, every participant discussed problems with how the consultations had been implemented and, in the UK, this critique also extended to the interpretation of the consultation:

> *… it seems to be very biased when I look at the study they performed and also the whole process now. You know, we’ve been invited to a conference, end of November. There’s a high-level conference -they call it, the EU - where they want to discuss the study and the whole consultation. And I think you can already read it in the title of the programme and in the title of the sessions where the way should lead. – NGO representative 3*

> *I mean it’s kind of [an] outrage. We are going to consult, we are going to consult… there’s no write up of that consultation, there’s no kind of big report that says these are the answers that we have got, these are the concerns, […] this is what people think where gene editing could help. It’s just like, yeah, we are going deregulate now. So how is that […] gaining public trust? – NGO representative 1*

The most common criticism of the consultation process from research and industry association representatives in both the EU and UK was “copy-paste” opinions: it was suggested that NGOs and certain political parties had asked members to provide email addresses so that pre-formed, template responses could be “copy-pasted” *pro-forma* on their behalf.

> *… so the green [parties] and NGOs who are circumventing the proper registration process… normally if you contribute you register with your email address then you get an email, you confirm with the link, so that it’s clear that it’s an individual with an email address that is contributing but this app just circumvented this process so you could fill in the app, and then immediately this contribution was showing up on the Commission’s website without proper verification of the email address, and the rules for those kind of consultations are that an individual can only provide one contribution. – Industry association representative 1*

On the other hand, UK-based NGO representatives we spoke to complained that the public comments weren’t properly considered in the decision-making process. The language used in the UK consultation was considered by UK participants to be too scientific or complex for lay people to understand.

One industry representative suggested that more open-ended, dialogic discussions would be a more effective way to feed public concerns into the consultation than is the current practice:

> *There will be a feeling among farming organisations colleagues that some of the environmental NGOs maybe just get up a standard letter that they get older members and supporters to stick their name on. And actually, when the civil servants are going through it they’ll discount a lot of that, so really, what’s the point? So maybe something deliberative, like a citizens’ jury type thing where you’re getting a group of citizens, people, you’ve got focus group deliberative exploration of the topic rather than a binary yes/no answer. – Industry association representative 4*

The length of active parliamentary rule making and the pace of the European Commission in clarifying the status of NGTs was also called into question by an EU industry association representative:

> *… we have the next European elections in 2024, and if you look at the policy processes [we] might end up in a situation that we will not have something in place until 2024. And then again we will only be able to, let’s say, reopen the discussion in 2025, 2026, so then it’s kind of already going into the end of [the decade]. – Industry association representative 1*

The ongoing influence of large multi-national companies – corporate interests in general – and their outsized role in the development of, at first, GMOs, and more recently NGTs, was treated with suspicion by NGO representatives. Beyond these concerns, though, many participants, regardless of their groups’ affiliations, mentioned that they were involved in collaborations and attempts to bring multiple stakeholders together to discuss NGTs:

> *… we looked for an agreement with the NGO’s involving the organic sector and we understand that [they] will never accept cisgenesis as non-GM technique*.*” – Industry association representative 2*

> *… another way of communicating could be what we have done in [country]… it was an open dialogue forum. So, face-to-face. Where we invited different stakeholders and the public and then had invited talks and discussion rounds. – Biotechnology research representative 2*

Many other elements of the European agri-food system, such as individual member states, the wider public and supermarkets, were named as influential by participants, who noted that professional or personal connections with decision makers and lobbying was (or was assumed to be) crucial for crop biotechnology policymaking.

## Discussion

Our data show that the stance of different stakeholder groups towards biotechnology remains rooted in its perceived risks, beginning with basic biology but extending to socio-economic considerations. These stances hold across both the European Union and United Kingdom. The two broad camps of opinion that formed around the use of first-generation, transgenic techniques have remained in place with respect to NGTs (see Macnaghten & Habets, 2020). These “lines” were already drawn by 2018 in advance of the ECJs decision on directed mutagenesis and the divisions were, as Poort et al. (2022) suggest, predictable. These two groups could be described as separate epistemic communities (see Haas et al., 1992) within Europe that frame problems – and solutions to those problems – in different terms, informed by different value systems and both seeking to influence policy according to what Lang and Barling (2012) view as competing ‘paradigms’ (see also Mampuys, 2022; Wynne, 2002). The use of biotechnology is one of several overlapping points of contention between groups with contrasting visions for the future of agriculture, environment and food.

Our findings reinforce existing research that perceptions of biotechnology risk vary considerably between stakeholders, making it difficult to reach agreement on acceptable levels of risk (Bouchaut & Asveld, 2020). Scepticism about the potential benefits of gene-edited organisms is more clearly articulated than the specific risks such products pose, though, which often consigned to vaguer notions of “unintended consequences” (echoing Wynne’s (2002) notion of ‘informal ambiguity’). A particular concern for those opposed to more lenient regulation for NGTs was entrenching corporate power in the food system through intellectual property rights in seed production – this stands in contrast to the gene editing imaginaries but forward by its proponents who see potential for the democratisation of agricultural biotechnologies through the use of NGTs (Bain et al., 2020). Coupled with the perception that so few of the systemic issues critics point to as problematic about first-generation techniques have changed (Montenegro de Wit, 2020), the “broken promises” of first-generation techniques have been inherited by NGTs even before their commercialisation in Europe (see also Kuiken et al., 2021). The legacy of first-generation biotechnology ensures that the impasse described by Macnaghten (2020) remains fixed, at least at the stakeholder level.

There were also concerns that scientific risk assessment, focussed on human and environmental health, is viewed by scientists and policymakers as a value-natural arena, rather than one that privileges expert knowledge and involves decisions that ultimately reflect the values of the members of decision-making groups (i.e. the ‘technocratic model’ (Scott, 2021)).

It was suggested that risk assessment could also include socio-economic assessment, thereby widening the pool of factors considered relevant to the regulation of specific products or technologies, including potential benefits (Bogner & Torgersen, 2018); this is consistent with Macnaghten & Habet’s (2020) framework and Kuiken et al’s (2021) suggestion that RRI be brought into the risk assessment of gene-edited crops in order to align NGT governance with the principles of responsible research. Existing research suggest that this process could involve a tiered system of case-by-case assessment for new applications, whereby those applications posing a greater risk could be subject to stakeholder consultations at the start of the development process and results fed back to developers, as proposed by the Norwegian Biotechnology Advisory Board (Bratlie et al., 2019). A number of recent European and North American public surveys and engagement activities demonstrate a continuum of biotechnology acceptance, from more acceptable purposes (human and crop disease resistance) to less (animal productivity or cosmetic interventions) (Busch et al., 2022; Mil et al., 2017). And, whilst between-country differences remain important when considering biotechnology acceptance (see Marette et al., 2021), the inclusion of means to weigh benefit against risk could embed *responsiveness*, a core principle of RRI, into the assessment process. On the basis of our findings, however, complications and potential criticism of this approach could stem from whether breeders have introduced more changes than a single point mutation (and whether these need be assessed) and opposition to *any* extra regulation in instances where these changes could have been achieved through conventional breeding.

Efforts to de-regulate NGTs is seen as a “backdoor” to de-regulation of transgenic crops (a concern attested to in Green European reports) (The Greens/EFA in the European Parliament, 2021). Despite recognition that plant breeding should align with societal needs (see Baekelandt & Parry, 2023) specific crop improvement aims, such as improving photosynthesis, could be judged superfluous where improving other outcomes – extreme poverty, for example – are seen to offer viable alternative strategies to meet social need over crop improvement. Previous research has identified that stakeholders in the food and farming community prefer plant breeding solutions when a lack of suitable alternatives exist to particular problems they face (Stetkiewicz et al., 2023). Despite rates of relative yield improvement being below requirements for the 2050 global population (Hall & Richards, 2013), improvement strategies targeting yield alone could struggle to be seen optimal given the range of issues that determine food (in)security; minimising yield losses in the face of a warming climate may be less controversial and, for research groups focussing on abiotic stress in crops, there is scope for better communication about why a plant breeding approach might be preferable.

The debate over NGT regulation was framed by our participants in a very similar way in both the EU and UK, despite the differences in regulatory regime emerging in the post-Brexit policy environment – this includes concerns over how public consultations were implemented, which were seen by the two camps of opinion as necessary but cursory (but for different reasons). Public trust is often cited as an important factor for biotechnology governance (Gordon et al., 2021; Hartley et al., 2016) and the need for some form of public involvement was reiterated across sampled groups. Our data suggest, though, that weaknesses in the consultatory process may contribute to a diminished trust in biotechnology policymaking. The official filtering of “genuine” responses from copy-paste opinions may have also contributed to a sense that the findings of the consultations (i.e. popular rejection of loosening rules) have been ignored in favour of a pre-determined policy towards NGTs.

More open-ended, deliberative and dialogic approaches, as promoted by some of our participants, could permit deeper understanding of the issues and/or provide a basis for new kinds of risk assessment. The concept of “mini-publics”, engaged through initiatives like citizen juries and assemblies (Willis et al., 2022), could be used to develop – and democratise – frameworks for biotechnology governance, such as have been used in discussions on climate change in the Republic of Ireland (Devaney et al., 2020) and the UK (Climate Assembly UK, 2020). Defra itself recently published a review, conducted by the Social Science Expert Group, which identified a number of examples of public engagement used in governance issues from air quality to bovine tuberculosis (Defra, 2022). More dialogic approaches also avoid the need for filtering responses – contributions to such a process are equally valuable. In contrast to the present consultations, it should be made transparent how these contributions would feed into policy (see Blasimme, 2019).

Mobilising political support through personal interaction with politicians was stated as important by industry representatives (and was assumed to be effective for industry by non-governmental organisation representatives). Some authors have argued it is the re-politicisation of this debate that offers a means to break the impasse that has not yielded to scientific input, participatory activities or regulatory approaches (Mampuys, 2022; Scott, 2021). Mampuys (2022) notes that science, participation and regulatory approaches do not compel a particular course of action – alternatively, a political decision built less on consensus and instead on compromise could provide a way forward (not unlike what has occurred in the UK). This could conflict with the “deliberative turn” in many fields of sustainability, such as science and technology, climate and food, which emphasise more and not less participation and delegation of power, but it is clear that some form of political support is seen as necessary by EU stakeholders for more favourable governance.

There was only marginal agreement, amongst those in the EU who wanted more lenient legislation for SDN-1 and -2 modification, on the best legislative ‘path’ forward; van der Berg et al. (2021) found a preference amongst the Dutch plant breeding sector for an exemption in existing legislation. As when our participants discussed this option, this was predicated on it being faster and more feasible than other options (such as the primary legislation now being discussed in the UK). An example exists in the exemption that was made for mutagenesis techniques in the 2001/18/EC directive, given their history of safe use.

## Limitations

Our approach has several limitations. The first relates to our positionality within debates around the regulation of biotechnology due to the nature of the PhotoBoost project. As a project that involves the use of crop biotechnology, some NGO representatives were wary of being involved in our interviews; these participants stressed that their involvement in the project via interview did not in itself constitute validation or approval for biotechnology projects, as might be claimed, they feared, by biotechnology researchers. We have attempted to mitigate these concerns through the inclusion of this statement and fair reporting of the themes identified in our work.

The second is that, in seeking balance between those for and against more lenient regulation for NGTs, we may be reinforcing this division; however, despite the low numbers of stakeholders involved in the research, the quick saturation of themes, relative rigidity in stakeholders views on these issues over time and often public campaigning for particular legislation, suggests this division is reflective of current opinions amongst key stakeholders.

The fast-moving regulatory environment also meant we captured the views of individuals *just prior* to the formal adoption of the Precision Breeding Bill in the UK and ahead of planned proposals from the EU with respect to its own regulations on NGTs in summer 2023 (indeed, some participants considered the process of establishing new regulation rushed).

## Conclusion

The European Union and United Kingdom are both in the process of establishing a new legislative framework for new genomic techniques in crop and animal breeding – each has sought some level of consultation in this process, but stakeholders on both sides of the debate around NGT regulation see flaws in how the consultations were carried out. The debate itself remains at an impasse rooted in risk, which for some groups extends beyond biological risk, with various approaches failing to provide a system that is agreeable to the wider agri-food sector and which has in turn led to calls to re-politicise the discussion that would allow policymakers to work towards a solution. It was through relationships with policymakers that stakeholders perceived the most progress could be made in shaping biotechnology regulation.

Our research suggests that there are opportunities to better engage stakeholders and the public either in developing science and technology policy, such as through citizens assemblies, or perhaps for specific applications of new genomic techniques where economic, societal and environmental impacts can be weighed. As some authors have suggested, there may be little consensus on the best path forward and research may best be mobilised to support the re-politicisation of this important debate.

## Acknowledgements

We would like to thank all those who took part in this study for their time and contributions, their input has been extremely valuable to our research.

